# *In Vivo* Expression of an SCA27A-linked *FGF14* Mutation Results in Haploinsufficiency and Impaired Firing of Cerebellar Purkinje Neurons

**DOI:** 10.1101/2024.10.25.620253

**Authors:** Joseph L. Ransdell, Samuel P. Brown, Maolei Xiao, David M. Ornitz, Jeanne M. Nerbonne

**Affiliations:** Department of Medicine, Cardiovascular Division, Washington University School of Medicine, St. Louis, MO 63110; Department of Biology, Miami University, Oxford, OH 45056; Department of Developmental Biology, Washington University School of Medicine, St. Louis, MO 63110

**Keywords:** iFGF14, iFGF14^F145S^, fibroblast growth factor homologous factor, FHF4, spinocerebellar ataxia type 27, cerebellum, intrinsic excitability, voltage-gated sodium (Nav) channels

## Abstract

Autosomal dominant mutations in *FGF14*, which encodes intracellular fibroblast growth factor 14 (iFGF14), underlie spinocerebellar ataxia type 27A (SCA27A), a devastating multisystem disorder resulting in progressive deficits in motor coordination and cognitive function. Mice lacking iFGF14 (*Fgf14^-/-^*) exhibit similar phenotypes, which have been linked to iFGF14-mediated modulation of the voltage-gated sodium (Nav) channels that control the high frequency repetitive firing of Purkinje neurons, the main output neurons of the cerebellar cortex. To investigate the pathophysiological mechanisms underlying SCA27A, we developed a targeted knock-in strategy to introduce the first point mutation identified in *FGF14* into the mouse *Fgf14* locus (*Fgf14^F145S^*), we determined the impact of *in vivo* expression of the mutant *Fgf14^F145S^* allele on the motor performance of adult animals and on the firing properties of mature Purkinje neurons in acute cerebellar slices. Electrophysiological experiments revealed that repetitive firing rates are attenuated in adult *Fgf14^F145S/+^* cerebellar Purkinje neurons, attributed to a hyperpolarizing shift in the voltage-dependence of steady-state inactivation of Nav channels. More severe effects on firing properties and Nav channel inactivation were observed in homozygous *Fgf14^F145S/F145S^* Purkinje neurons. Interestingly, the electrophysiological phenotypes identified in adult *Fgf14^F145S/+^*and *Fgf14^F145S/F145S^* cerebellar Purkinje neurons mirror those observed in heterozygous *Fgf14^+/-^* and homozygous *Fgf14^-/-^*Purkinje neurons, respectively, suggesting that the mutation results in the loss of the iFGF14 protein. Western blot analysis of lysates from adult heterozygous *Fgf14^F145S/+^* and homozygous *Fgf14^F145S/F145S^*animals revealed reduced or undetectable, respectively, iFGF14 expression, supporting the hypothesis that the mutant allele results in loss of the iFGF14 protein and that haploinsufficiency underlies SCA27A neurological phenotypes.

## Introduction

Although sharing sequence homology with canonical members of the fibroblast growth factor (FGF) superfamily, the intracellular fibroblast growth factors, iFGF11-14, also called fibroblast growth factor homologous factors (FHFs), are distinct (Ornitz and Itoh, 2022; Pablo and Pitt, 2016). The iFGFs, for example, are expressed in adult tissues, as well as during development, and lack signal sequences for secretion (Goldfarb, 2005; Hartung et al., 1997; Olsen et al., 2003; Smallwood et al., 1996; Wang et al., 2000). Although there is evidence suggesting that the iFGFs can, under some circumstances, activate FGF receptors (Biadun et al., 2023; Sochacka et al., 2020), the primary physiological function identified to date for the iFGFs reflect binding to the intracellular C-terminal domains of voltage-gated sodium (Nav) channel pore-forming (α) subunits (Bosch et al., 2016; Goetz et al., 2009; Liu et al., 2003; Liu Cj et al., 2001; Lou et al., 2005; Wang et al., 2011; Wittmack et al., 2004) and modulation of the time- and voltage-dependent properties of Nav currents (Abrams et al., 2020; Angsutararux et al., 2023; Bosch et al., 2015; Chakouri et al., 2022; Goldfarb et al., 2007; Laezza et al., 2009; Lou et al., 2005; Park et al., 2016; Wittmack et al., 2004; Yan et al., 2014). The functional effects on Nav currents observed, however, depend on the specific iFGF and Nav α subunit that are co-expressed and also on the cellular environment (Di Re et al., 2017; Pablo and Pitt, 2017; Ransdell et al., 2024).

*In vivo*, iFGF14 localizes to the axon initial segments of hippocampal pyramidal neurons (Goldfarb et al., 2007; Laezza et al., 2007; Lou et al., 2005; Pablo and Pitt, 2017; Ransdell et al., 2024), as well as cerebellar Purkinje, granule, basket, and stellate neurons (Bosch et al., 2015; Xiao et al., 2013; Yan et al., 2014). Animals harboring targeted disruption of the *Fgf14* locus (*Fgf14^-/-^*) display severe locomotor and learning deficits (Wang et al., 2002; Wozniak et al., 2007). Consistent with the locomotor phenotype observed in *Fgf14^-/-^* animals (Wang et al., 2002), cellular electrophysiological studies demonstrated that *Fgf14^-/-^* cerebellar Purkinje neurons lack the spontaneous repetitive firing that is characteristic of wild type Purkinje neurons (Bosch et al., 2015; Shakkottai et al., 2009). Deficits in action potential firing were also observed with acute shRNA-mediated *in vivo* knockdown of *Fgf14* in adult Purkinje neurons, and were linked to a hyperpolarizing shift in the voltage-dependence of closed state inactivation of the transient Nav current (Bosch et al., 2015). It has also been reported that acute shRNA-mediated knockdown of *Fgf14* in isolated neonatal Purkinje neurons *in vitro* increased the rate of transient Nav current inactivation and reduced the resurgent Nav current (Yan et al., 2014), effects that are *not* observed in adult cerebellar Purkinje neurons with *Fgf14* knockdown or knockout (Bosch et al., 2015).

Shortly after the description of the *Fgf14^-/-^* mouse line (Wang et al., 2002), an autosomal dominant missense mutation in *FGF14*, resulting in a single (phenylalanine to serine) amino acid substitution in iFGF14, was identified in a large Dutch family presenting with ataxia and progressive cognitive decline (van Swieten et al., 2003). Several additional autosomal dominant mutations in *FGF14*, including frameshift (Choquet et al., 2015; Dalski et al., 2005), nonsense (Miura et al., 2019), and chromosomal translocations (Misceo et al., 2009; Shimojima et al., 2012) were subsequently identified, and these are now collectively classified as spinocerebellar ataxia type 27A (SCA27A). Interestingly, GAA repeat expansions in *FGF14* were recently identified as one of the most common genetic causes of late-onset ataxias referred to as SCA27B (Kartanou et al., 2024; Pellerin et al., 2024; Rafehi et al., 2023; Wilke et al., 2023).

Modeling studies suggested that the F145S mutation will result in unstable iFGF14 protein and therefore, loss of iFGF14 function (Olsen et al., 2003; van Swieten et al., 2003). Expression of the iFGF14-F145S mutant protein in neonatal rat hippocampal neurons *in vitro*, however, attenuated Nav current amplitudes and reduced neuronal excitability (Laezza et al., 2007). In additional experiments conducted in heterologous (HEK-293) cells, the iFGF14-F145S mutant protein was shown to function as a dominant-negative, preventing wild type iFGF14 from interacting with and modulating Nav currents (Laezza et al., 2007). To explore the mechanism(s) by which autosomal dominant mutations in *FGF14* affect *in vivo* neurological functioning, we developed a mouse model with targeted knock-in of the human *FGF14^F145S^* mutation at the *Fgf14* locus and determined the impact of the *in vivo* expression of the iFG14-F145S protein on Purkinje neuron excitability and cerebellar functioning directly. These experiments collectively reveal autosomal dominant impairments in *Fgf14^F145S/+^* animals are caused by haploinsufficiency.

## Materials and Methods

All reagents were obtained from Sigma-Aldrich unless otherwise indicated.

### Experimental Animals

All experiments involving animals were performed in accordance with the guidelines published in the National Institutes of Health *Guide for the Care and Use of Laboratory Animals.* All protocols were approved by the Washington University or Miami University Institutional Animal Care and Use Committees (IACUC). To construct an *in vivo* mouse model, we selected the first human mutation, FGF14^F145S^, identified (van Swieten et al., 2003), and mutated the codon for phenylalanine 145 (F145), as illustrated in Figure 1. F145 is based on the human FGF14-1a numbering system and corresponds to F149 in the mouse sequence. The F145 codon (TTT) is located in exon 4 and overlaps with a Dra1 restriction site (TTTAAA). The human mutation, TTT to TCT, replaces phenylalanine with serine (S) and destroys the Dra1 restriction site. The nucleotide sequence encoding F145 and the Dra1 restriction site is identical between mouse and human. The targeting vector depicted in Figure 1A along with homologous recombination methods using the SCC#10 ES cell line (ATCC SCRC-1020, RRID:CVCL_C259) were used to generate the F145 *Fgf14* mutation. Clones that showed both 5’ and 3’ homologous recombination were screened for loss of the Dra1 restriction site. Wild type DNA, amplified with primers flanking exon 4, were expected to cut with Dra1, while targeted DNA that contained the engineered mutation were expected not to cut with Dra1. Cell lines that contained the targeted mutation were then karyotyped. Cells with a normal, 40, XY karyotype that contained the F145S mutation were injected into C57BL/6 blastocysts by the Washington University Mouse Genetics Core. Resulting chimeric mice were bred and tested for germline transmission. Mice containing a targeted allele were bred to mice expressing Cre recombinase in the germline. This resulted in the excision of the LoxP-neo-LoxP cassette, leaving behind only a single loxP site (Figure 1A).

**Figure 1.**
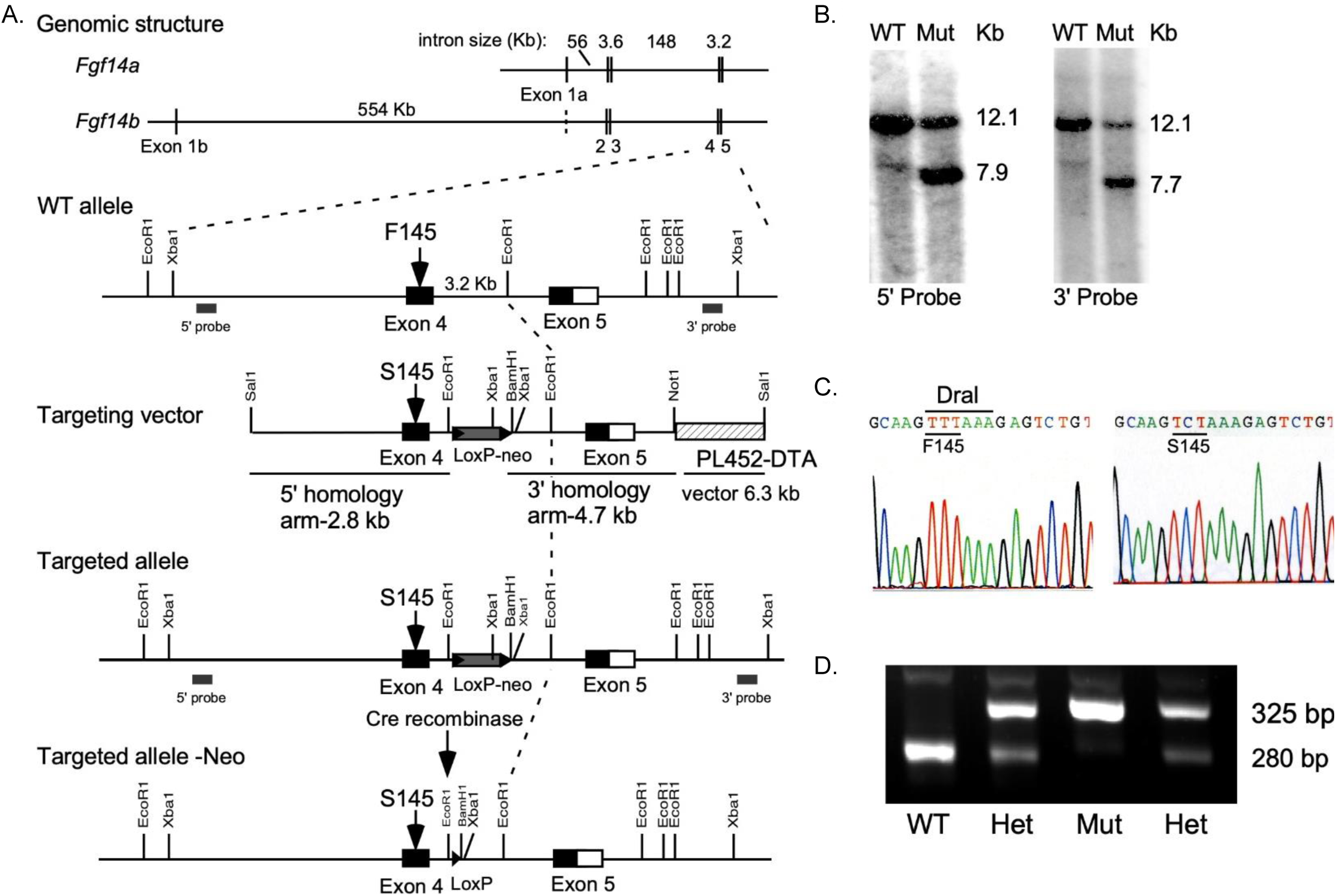
Mouse model of SCA27: knock-in expression of the iFGF14^F145S^ human mutation. (**A**) Schematic of the wild type *Fgf14* gene structure and targeting strategy to introduce the F145S mutation. Top, genomic structure showing exons 1b-5. Below, the targeting construct with a *neo* cassette and flanking *loxP* sequences, the mutant allele produced by homologous recombination, and the mutant allele with *neo* deletion mediated by Cre. (**B**) Southern blot analysis (Xba1 digest) with 5’ and 3’ probes (labeled in **A**) allowed the identification of the mutant and wild type alleles (after *neo* deletion). (**C**) Sanger sequencing chromatograms of genomic PCR products confirmed the wild type F145 and homozygous S145 sequences. (**D**) Representative gel illustrating the PCR genotyping results of littermates from a *Fgf14^F145S/+^* x *Fgf14^F145S/+^* cross with primers flanking the *loxP* site. Lane 1 shows the wild type allele (280 bp); lanes 2 and 4 show the heterozygote mutant (*Fgf14^F145S/+^*), wild type (280 bp), mutant (325 bp) (after *neo* deletion), and lane 3 shows the homozygous mutant (*Fgf14^F145S/F145S^*) (325 bp).

Animals harboring a targeted disruption of the *Fgf14* locus (*Fgf14^-/-^*), described in Wang et al. (2002), were generated by crossing heterozygous *Fgf14^+/-^* mice, congenic in the C57BL/6 background; genotypes were confirmed by PCR. Homozygous *Fgf14^F145S/F145S^*knock-in animals were generated by crossing heterozygous *Fgf14^F145S/+^*(heterozygotes) in the C57BL/6 background. Genotypes were confirmed by PCR screening. Male and female mice (P12 to 8 weeks-old) of the genotypes (*Fgf14^+/-^, Fgf14^-/-^*, *Fgf14^F145S/+^*, *Fgf14^F145S/F145S^*), along with wild type controls, were used in the experiments completed here.

### Elevated balance beam

The elevated balance beam assay was performed using previously described methods (Ransdell et al., 2017). Briefly, animals were trained for three consecutive days, followed by five consecutive days of testing. During each training day, animals were allowed to explore the Plexiglas platform and the enclosed box for two minutes on each surface. Subsequently, during training, animals were placed on the Plexiglas platform and allowed to transverse an 11 mm-wide beam into an enclosed box. After 3 days of training, animals began five consecutive days of testing. During testing days, animals (individually) crossed the 11 mm-wide beam (from the platform to the enclosed box) and both the time to cross and the number of hind-limb foot-slips during crossing were recorded by the experimenter. During all tests, the experimenter was blinded to animals’ genotypes.

### Preparation of acute brain slices

Acute (350 μM) parasagittal cerebellar slices were prepared from (5-8 week-old) animals using previously described methods (Bosch et al., 2015; Ransdell et al., 2017). Briefly, animals were anaesthetized with 1.25% Avertin and perfused transcardially with ice-cold cutting solution that contained (in mM): 240 sucrose, 2.5 KCl, 1.25 NaH_2_PO_4_, 25 NaHCO_3_. 0.5 CaCl_2_, and 7 MgCl_2_ saturated with 95% O_2_/5% CO_2_. Following perfusion, animals were rapidly decapitated and the brain of each animal was removed and cut parasagittally in ice-cold cutting solution on a Leica VT1000S vibratome (Leica Microsystems Inc. Buffalo Grove, IL, USA). Parasagittal cerebellar sections were incubated in warmed (34°C) artificial cerebrospinal fluid (ACSF) containing (in mM): 125 NaCl, 2.5 KCl, 1.25 NaH_2_PO_4_, 25 NaHCO_3_, 2 CaCl_2_, 1 MgCl_2_, and 25 dextrose, ∼310 mOsmol/L, saturated with 95% O_2_/5% CO_2_, for 35 minutes and an additional 25 minutes – 4 hours at room temperature (25°C) prior to electrophysiolgoical recordings.

### Isolation of neonatal cerebellar Purkinje neurons

P12-P15 wild type and *Fgf14^-/-^* animals were anesthetized with 1.25% Avertin and brains were rapidly removed and placed in ice-cold isolation medium containing (in mM): 82 Na_2_SO_4_, 30 K_2_SO_4_, 5 MgCl_2_, 10 HEPES, 10 glucose, and 0.001% phenol red (at pH 7.4). Using a scalpel, the cerebellum was removed, minced into small chunks and incubated in isolation medium containing 3 mg/mL protease XXIV at 33°C for 10-15 minutes. Following this incubation period, the tissue pieces were washed with enzyme-free isolation medium containing 1 mg/mL bovine serum albumin and 1 mg/mL trypsin inhibitor. After being transferred to oxygenated ACSF (22–23°C), the tissue pieces were triturated with a fire-polished glass pipette. An aliquot of the cell suspension was placed on a coverslip in the recording chamber and superfused with fresh ACSF (0.5 mL / min) saturated with 95% O2/5% CO2 for 25 minutes before beginning electrophysiological recordings.

### Electrophysiological recordings

Whole-cell current-clamp and voltage-clamp recordings were obtained from visually identified cerebellar Purkinje neurons using differential interference contrast optics. During recordings from cells in parasagittal cerebellar slices, warmed ACSF (34 ± 1°C) saturated with 95% O_2_/5% CO_2_ was continuously perfused. Data were collected using a Multiclamp 700B patch clamp amplifier interfaced with a Digidata 1332 and pCLAMP 10 software (Axon Instruments, Union City, CA, USA). In all experiments, tip potentials were zeroed prior to forming giga-ohm membrane-pipette seals. Pipette capacitances were compensated using the pCLAMP software. Signals were acquired at 50-100 kHz and filtered at 10 kHz prior to digitization and storage.

For current-clamp recordings, pipettes contained (in mM): 144 K-gluconate, 0.2 EGTA, 3 MgCl_2_,10 HEPES, 4 MgATP, 0.5 NaGTP, PH 7.3, ∼ 300 mosomol/L. Spontaneous firing was recorded for 60 seconds and analyzed during the last 30 seconds. Evoked repetitive firing was measured during 0.5 s depolarizing current injections of varying amplitudes that were administered immediately after a -0.5 nA prepulse. For voltage-clamp recordings, pipettes contained (in mM): 110 CsCl, 15 TEA-Cl, 5 4AP, 1 CaCl_2_, 2 MgCl_2_, 10 EGTA, 4 Na_2_-ATP and 10 HEPES, pH 7.25 ∼300 mOsmol/L. Recording pipettes had resistances of 2-4 MΩ. Series resistances were compensated ≥80%, and the voltage errors resulting from the uncompensated resistances were always ≤ 4mV and were not corrected. To isolate and measure Nav currents in Purkinje neurons in acute cerebellar slices, after formation of a gigaseal, the superfusing ACSF was switched to a low Na^+^ ACSF containing (in mM): 25 NaCl, 100 TEA-Cl, 2.5 KCl, 1.25 NaH_2_PO_4_, 25 NaHCO_3_, 2 CaCl_2_, 1 MgCl_2_, 0.1 CdCl_2_, and 25 dextrose, ∼310 mOsmol/L, saturated with 95% O_2_/5% CO_2_. Voltage-clamp protocols to isolate Nav currents were tested using a strategy outlined in Milescu et al. (2010), and optimized to minimize space-clamp errors and enable reliable measurements of Nav currents in Purkinje neurons. To determine the voltage-dependence of activation of the Nav currents, a three-step voltage-clamp protocol was used: a depolarizing pre-pulse (0 mV, 5 ms), to inactivate the Nav currents in the distal neurites, was delivered first, followed by a brief (2.5 ms) hyperpolarizing voltage step to -60 mV, to allow recovery of the Nav currents in the soma and proximal neurites and a second depolarizing pulse (for 8ms) was used to measure (appropriately clamped) Nav currents at different test potentials. Peak Nav conductance (at each potential) was calculated (using E_reversal_ = +43 mV) and normalized to the maximal Nav conductance. The mean ± SEM normalized Nav conductances were plotted as a function of the test potential, and fitted with a single (Equation 1) Boltzmann function shown below:

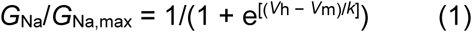

Where *V*_h_ is the membrane potential of half-maximal activation and *k* is the slope factor.

To determine the voltage-dependence of steady-state inactivation of the Nav currents, a three step voltage-clamp protocol was also used. Each cell was first depolarized to 0 mV for 5 ms to inactivate the Nav currents, and subsequently hyperpolarized briefly (2.5 ms) to various conditioning voltages to allow recovery of Nav channels from inactivation and, finally, depolarized to -20 mV for 8 ms. The peak transient Nav current amplitudes evoked during this final step (to - 20 mV) were measured and normalized to the current amplitude evoked (at -20 mV) from a -120 mV conditioning voltage step (in the same cell). Mean ± SEM normalized Nav currents were plotted as a function of the conditioning voltage. The voltage-dependence of Nav current steady-state inactivation were plotted and fit with single Boltzmann functions (Equation 2).

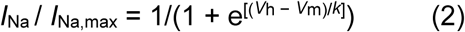

Where *V*_h_ is the membrane potential of half-maximal inactivation and *k* is the slope factor.

### Analysis of Electrophysiological Data

Data were analyzed using ClampFit (Molecular Devices), MATLAB (Mathworks), Microsoft Excel, and Prism (GraphPad Software Inc.). Input resistances (*R*_in_) were determined from the change in membrane potential produced by a 20 pA hyperpolarizing current injection from the resting potential. Percentages of time firing were calculated using a custom MATLAB script in which 30 seconds of spontaneous (current-clamp) activity was analyzed and the elapsed times between action potentials (inter-spike interval) were categorized as ‘firing’ if less than or equal to 50 ms. If the interspike-intervals were greater than 50 ms, the elapsed times between action potentials were characterized as ‘silent’. In this analysis, sodium-mediated spikes were recognized (and differentiated from calcium spikes) as having maximal rising phases of greater than or equal to 100 mV/ms. These values were based on previous experiments in our lab (not shown) in which TTX (which blocks Nav channels) and cadmium (which blocks calcium channels) were used to determine that Purkinje neuron calcium-mediated spikes, measured after applying 1 µM TTX, have maximum rising dV/dt values that are < 100 mV/ms and sodium mediated spikes have maximum rising dV/dt values that are > 100 mV/ms.

Averaged and normalized data are presented as means ± SEM. Statistical analyses were performed using Student’s paired or unpaired t-tests or two-way analysis of variance (ANOVA) as identified in the text and/or the figure legends.

### Western Blot Analysis

Lysates were prepared from adult (> 8 week-old) animals in 20 mM HEPES + 150 mM NaCl buffer with 0.5% CHAPS and a protease inhibitor tablet (Roche) using established methods (Brunet et al., 2004; Norris and Nerbonne, 2010), fractionated on SDS-PAGE (12%) gels, transferred to polyvinylidene fluoride (PVDF) membranes (Biorad) and probed for iFGF14 expression using a rabbit polyclonal anti-iFGF14 antiserum (1:1,000; Covance Laboratories), validated as described previously (Bosch et al., 2016).

## Results

### Development of a SCA27A mouse model expressing the FGF14^F145S^ mutation

A gene-targeting construct for knock-in expression of the human missense *FGF14^F145S^* mutation was developed site specifically (see Materials and Methods and Figure 1A). Southern blot analysis (Xba1 digest) with 5’ and 3’ probes (locations labeled in Figure 1A) allowed the identification of the mutant and wild type alleles in two independent embryonic stem cell clones (Figure 1B) and Sanger sequencing of the genomic PCR products confirmed the presence of the wild type (WT), the homozygous S145 mutation, and the heterozygous F145/S145 sequences (Figure 1C). Animals (*Fgf14^F145S/+^*) heterozygous for the mutant allele were crossed to generate offspring that were heterozygous (*Fgf14^F145S/+^*) or homozygous (*Fgf14^F145S/F145S^*) for the *Fgf14* mutation. Offspring (from this and subsequent crosses) were screened by PCR using the primers given in the legend to Figure 1 and representative results are illustrated in Figure 1D. Heterozygous *Fgf14^F145S/+^*animals were identified by the presence of bands corresponding to both the wild type (280 base pairs) and the mutant (325 base pairs) alleles and homozygous *Fgf14^F145S/F145S^* animals were identified by the presence of only the mutant (325 base pairs) allele (Figure 1D).

### Sustained repetitive firing is impaired in *Fgf14^F145S/+^* Purkinje neurons

To determine how heterozygous expression of the iFGF14^F145S^ human mutation affects the intrinsic excitability of cerebellar Purkinje neurons, we obtained whole-cell current clamp recordings from *Fgf14^F145S/+^*Purkinje neurons in acute parasagittal cerebellar slices prepared from adult (5-8 week old) animals at (34 ± 1°C). These experiments revealed that the high frequency repetitive firing, which is characteristic of wild type Purkinje neurons in acute cerebellar slices (Häusser and Clark, 1997), was impaired in *Fgf14^F145S/+^* Purkinje neurons. In contrast to the sustained repetitive spike activity typically measured in wild type Purkinje neurons (Figure 2A), *Fgf14^F145S/+^* Purkinje neurons fired in lengthy bursts of sodium- and calcium-mediated spikes, and these bursts were punctuated by periods of quiescence (Figure 2B). Only rarely (3/26 cells), did *Fgf14^F145S/+^* Purkinje neurons fire repetitively for the entire 30 second analysis period.

**Figure 2.**
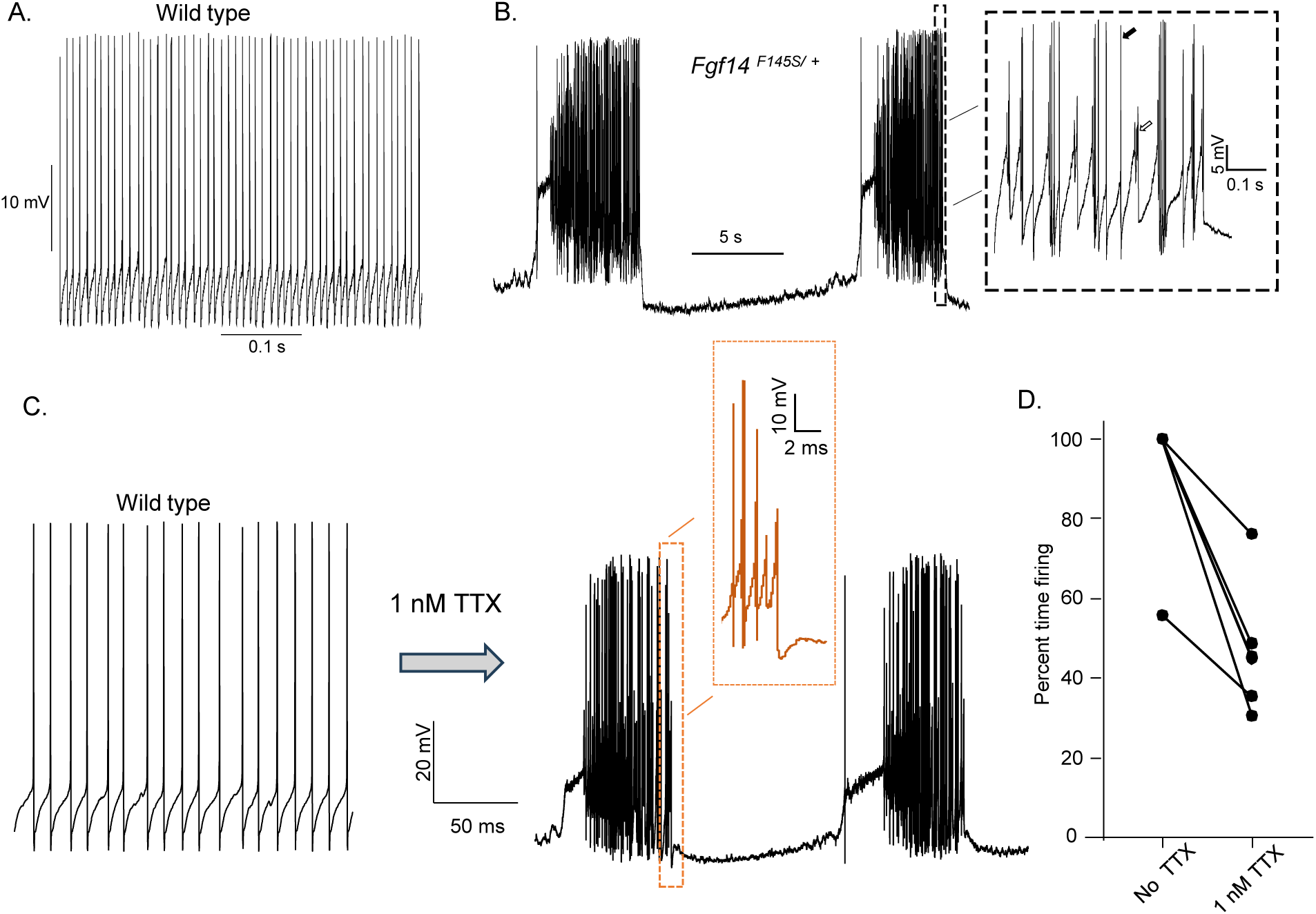
Sustained High Frequency Firing is Impaired in iFGF14^F145S^ Cerebellar Purkinje Neurons and Mimicked by Treatment of wild type Cells with TTX. (**A-B**). Representative recordings of spontaneous firing obtained from adult Purkinje neurons in acute cerebellar slices from (**A**) wild type and **(B**) *Fgf14^F145S/+^*animals are shown. The Inset in panel **B** highlights the sodium- (filled arrow) and calcium- (unfilled arrow) mediated spikes in the representative *Fgf14^F145S/+^* Purkinje neuron (see Methods). (**C**) Representative recordings from an adult wild type Purkinje neuron before and after the addition of a 1 nM TTX to the perfusing ACSF converts sustained repetitive firing into biphasic periods of quiescence and firing, similar to the biphasic firing pattern observed (**B**) in *Fgf14^F145S/+^* Purkinje neurons. (**D**) Addition of 1 nM TTX to wild type Purkinje neurons markedly (*P* = .002, paired Student’s t-test, n = 6) reduced the mean (± SEM) percentage of time of repetitive action potential firing measured during 30 second current-clamp recordings.

We hypothesized the intermittent spontaneous activity recorded in *Fgf14^F145S/+^* Purkinje neurons may reflect reduced Nav channel availability in these cells. To explore this hypothesis, we applied a low concentration (1 nM) of TTX that has previously been demonstrated to attenuate, but not eliminate, Nav currents in Purkinje neurons (Khaliq and Raman, 2006). As illustrated in Figure 2C, the characteristic high frequency repetitive firing of a wild type Purkinje neuron is altered dramatically on addition of 1 nM TTX. In the presence of 1 nM TTX, Purkinje cells fire in prolonged bursts of action potentials, which also included slower, calcium-mediated spikes; firing behavior that is similar to the spontaneous firing recorded in *Fgf14^F145S/+^* Purkinje neurons (Figure 2B). To quantify firing rates, we calculated the mean ± SEM percentage of time wild type Purkinje neurons fired sodium-mediated action potentials during a 30 second whole-cell recording period, before and after exposure to 1 nM TTX. These analyses revealed markedly (*P* = .002) reduced sodium-mediated spike activity in the presence of 1 nM TTX. The loss of sustained repetitive firing and the conversion to calcium-mediated spikes in the presence of 1 nM TTX suggests this firing phenotype reflects reduced Nav channel availability.

### Effect of *Fgf14^F145S^* on Purkinje neuron repetitive firing phenocopies *Fgf14* deletion

Additional experiments revealed that the spontaneous firing properties of Purkinje neurons lacking a single *Fgf14* allele (*Fgf14^+/-^*) is very similar (Figure 3B) to the spontaneous firing observed in *Fgf14^F145S/+^* Purkinje neurons (Figure 2B), with prolonged bursts of sodium- and calcium-mediated spikes that are interrupted by periods of quiescence. We also measured spontaneous firing in adult Purkinje neurons with homozygous knock-in of the *Fgf14^F145S^* mutant allele (*Fgf14^F145S/F145S^*) and found these cells are mostly silent (Figure 3C) in stark contrast to the maintained high frequency repetitive firing that is typically observed in wild type Purkinje neurons (Figure 3A): 11/13 *Fgf14^F145S/F145S^*Purkinje neurons were found to be silent during the 30 second recording period. This result is similar to recordings from *Fgf14^-/-^* Purkinje neurons in which the vast majority (14/16) of cells were also found to be silent (Figure 3D). These spontaneous firing phenotypes, which are presented as representative records in Figures 2A-B and 3A-C, were quantified by calculating the proportion of time (during a 30 second current-clamp recording) that each Purkinje neuron fired repetitive, sodium-mediated action potentials. This is the same analysis that we conducted on TTX-exposed Purkinje neurons presented in Figure 2D. The duration between action potentials was categorized as repetitive if the inter-spike intervals were ≤ 50 ms, and spikes were determined to be sodium-mediated if the rising phases had a peak dV/dt value of ≥100 mV/ms (see Methods). As the representative records suggest, the mean ± SEM percentages of time Purkinje neurons from *Fgf14^F145S/+^*(33 ± 7%) and *Fgf14^+/-^* (36 ± 11%) animals fire repetitively are similar (Figure 3D), and intermediate to the mean ± SEM percentages of time that *Fgf14^F145S/F145S^* (4 ± 3%), *Fgf14^-/-^* (4 ± 4%), and wild type (75 ± 5%) Purkinje neurons repetitively fire sodium-mediated action potentials (Figure 3D).

**Figure 3.**
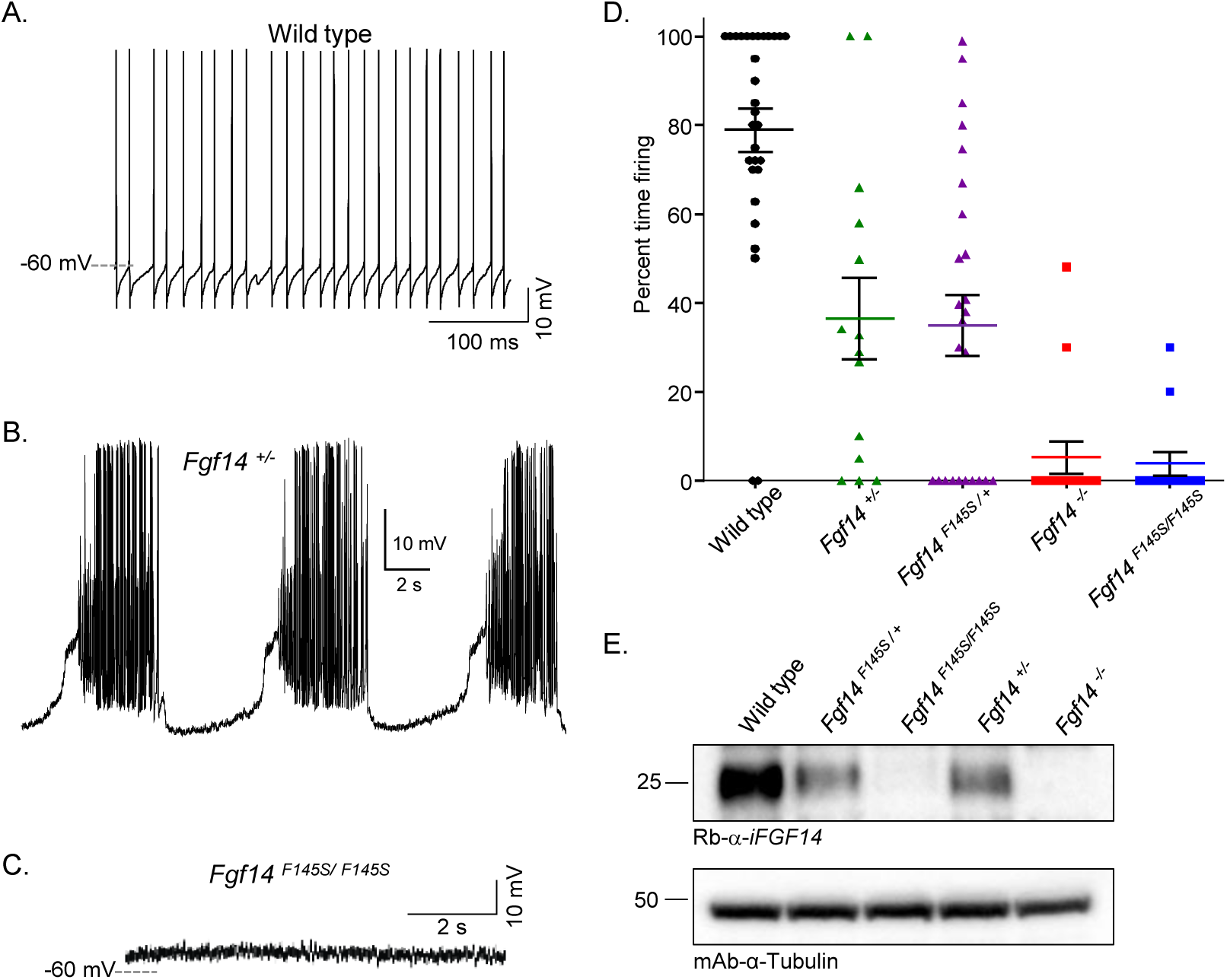
Repetitive firing properties of *Fgf14^F145S/+^* and *Fgf14^F145S/F145S^*Purkinje neurons phenocopy the firing properties of *Fgf14^+/-^* and *Fgf14^-/-^* Purkinje neurons. (**A-C**). Representative recordings of spontaneous firing from Purkinje neurons in acute cerebellar slices from adult wild type (**A**), *Fgf14^+/-^* (**B**), and *Fgf14^F145S/F145S^* (**C**) animals are shown. (**D**). The mean ± SEM percentages of time firing (during 30 second current-clamp recordings), determined for each genotype, are plotted and reveal that *Fgf14^+/-^* and *Fgf14^F145S/+^* Purkinje neurons, lacking one copy of the wild type *Fgf14* allele, fire repetitive sodium-mediated action potentials much less than wild type Purkinje neurons. (**E**). Representative Western blots of cerebellar lysates prepared from adult wild type, *Fgf14^F145S/+^, Fgf14^F145S/F145S^*, *Fgf14^+/-^*, and *Fgf14^-/-^*animals probed with a rabbit polyclonal iFGF14 antiserum and with mouse monoclonal anti-tubulin antibody (loading control) reveal reduced iFGF14 expression in cerebellar tissues from animals with targeted knock-in of the *Fgf14^F145S^*mutant allele. Full Western blot gel images are shown in Extended Data Figure 3-1.

### Knock-in of the *Fgf14^F145S^* mutant allele reduces iFGF14 protein

Results from the current clamp studies described above reveal that knock-in expression of the *Fgf14^F145S^* mutant allele has effects similar to targeted *Fgf14* deletion. To determine if the *Fgf14^F145S^*mutation results in loss of the iFGF14 protein, cerebellar tissue lysates from *Fgf14^F145S/F145S^* and *Fgf14^F145S/+^* animals were fractionated, probed with an anti-iFGF14 antiserum, and compared directly to cerebellar tissue lysates from wild type animals and from animals with targeted deletion of one (*Fgf14^+/-^*) or both (*Fgf14^-/-^*) *Fgf14* alleles; a representative Western blot is presented in Figure 3E. As is evident, these analyses revealed that the anti-iFGF14 signals from heterozygous knock-in (*Fgf14^F145S/+^*) and knock-out (*Fgf14^+/-^*) tissue lysates are lower than the anti-iFGF14 signal in wild type cerebellar tissue lysates (Figure 3E). In addition, similar to *Fgf14^-/-^* tissue lysates, the iFGF14 protein is undetectable in cerebellar tissue lysates prepared from *Fgf14^F145S/F145S^* animals indicating that *in vivo* expression of the *Fgf14^F145S^* missense mutation results in loss of the iFGF14 protein.

### Similar to *Fgf14* deletion, knock-in of *Fgf14^F145S^* affects Nav current inactivation

To explore the molecular mechanisms underlying the repetitive firing phenotypes observed in *Fgf14^F145S/+^* (Figure 2) and *Fgf14^F145S/F145S^* (Figure 3) cerebellar Purkinje neurons, we examined the voltage-dependences of activation and inactivation of the fast-transient Nav currents (I_NaT_). To accurately measure Nav currents from mature Purkinje neurons in acute brain slices, and avoid errors caused by inadequate space-clamp that are typical in experiments involving cells with intact neurites, voltage-clamp experiments were performed (see Materials and Methods) with reduced extracellular Na^+^ and using a voltage-clamp strategy developed by Milescu et al. (2010) (see Materials and Methods). Briefly, cells were first depolarized to activate and inactivate the Nav channels in the soma and distal neurites, followed by brief hyperpolarization to allow recovery of Nav channels in/near the soma, and subsequent depolarization to various test potentials to measure Nav current amplitudes and the voltage-dependent properties of recovered Nav channels at different test potentials. Figure 4A depicts the voltage-clamp protocol (lower) and representative Nav current records (upper) used to examine the voltage-dependence of Nav current steady-state inactivation. Peak current amplitudes, measured at the -20 mV test potential, were normalized to the current amplitude after a conditioning voltage-step to -120 mV. The mean ± SEM normalized Nav current amplitudes are plotted as a function of the conditioning membrane potential (Figure 4B) and Boltzmann fits of these plots (see Materials and Methods) reveal that 50% of the Nav channels are inactivated (V_1/2_) in *Fgf14^F145S/F145S^* (V_1/2_ = -71 mV, n = 10) and *Fgf14^F145S/+^* (V_1/2_ = -65 mV, n = 8) Purkinje neurons at voltages that are significantly (*P* ≤ .0001, two-way ANOVA) different from one another, and from wild type (V_1/2_ = -58 mV, n = 11) Purkinje neurons. The slopes of the fits to the steady-state inactivation data (Fig. 4B) in *Fgf14^F145S/F145S^*, *Fgf14^F145S/+^*, and wild type Purkinje neurons are 10.5 ± 0.6, 9.9 ± 0.7, and 11 ± 0.9, respectively. Conversely, plots of normalized Nav conductance against activation voltage are not significantly different in wild type, *Fgf14^F145S/+^*, and *Fgf14^F145S/F145S^* Purkinje neurons (Figure 4C).

**Figure 4.**
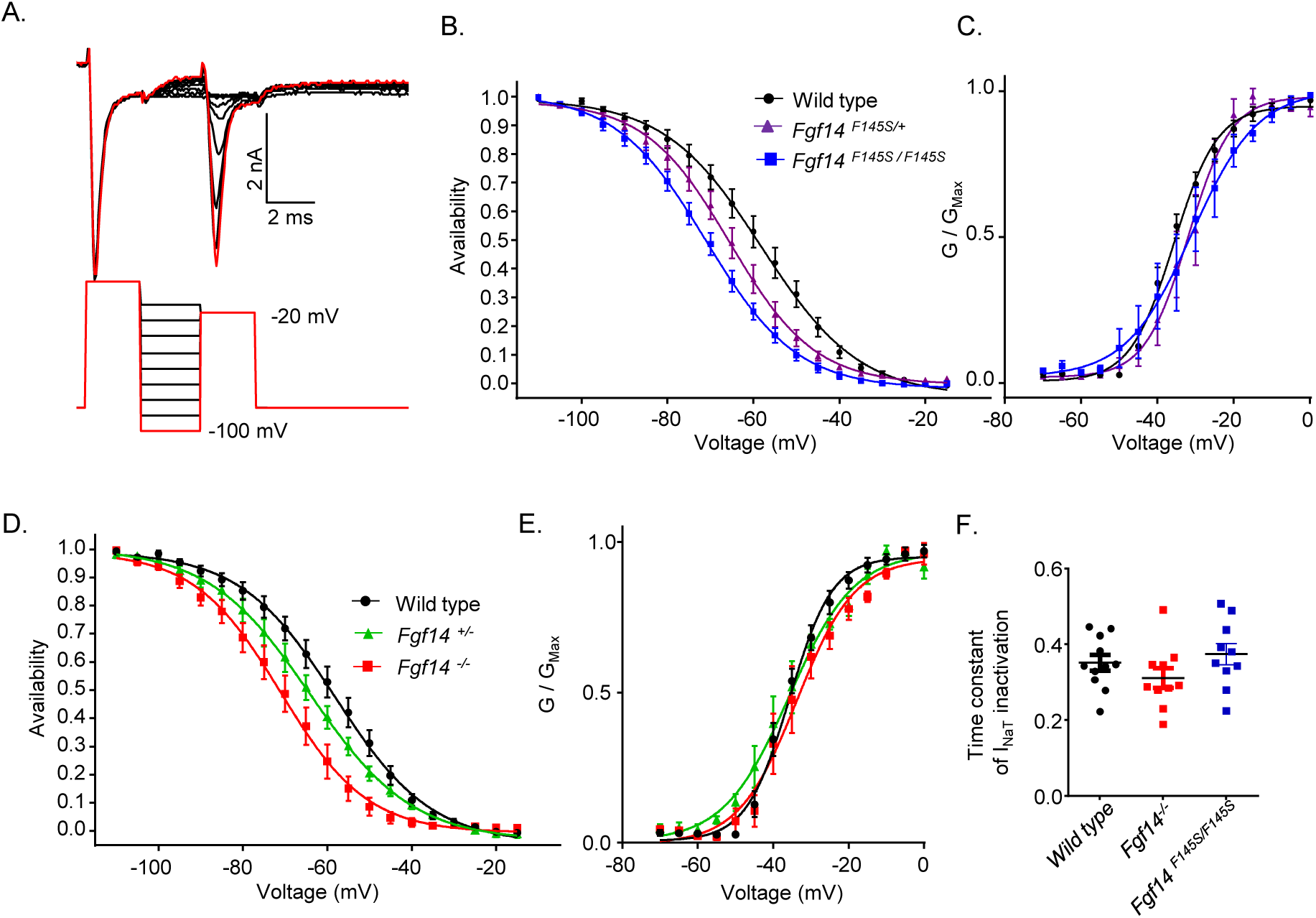
Expression of *Fgf14^F145S^* causes changes in Nav currents in Purkinje neurons similar to targeted *Fgf14* deletion. (**A**). To analyze the voltage-dependence of steady-state inactivation of I_NaT_ in Purkinje neurons across genotypes, peak I_NaT_ amplitudes evoked at -20 mV from a 2.5 ms voltage step to various conditioning potentials (-120 mV to -10 mV) were measured and normalized to the peak I_NaT_ measured from the −120 mV conditioning voltage step in the same cell; the voltage-clamp paradigm is illustrated below the representative current records. (**B**). Mean ± SEM normalized peak I_NaT_ amplitudes in wild type (*black*; n = 11), *Fgf14^F145S/+^* (*purple*, n = 8) and *Fgf14^F145S/F145S^* (*blue*, n = 10) Purkinje neurons are plotted as a function of the conditioning voltage and fitted with first order Boltzmann functions. (**C**). To determine the voltage-dependences of I_NaT_ activation, the mean ± SEM normalized peak I_NaT_ conductances calculated (as described in Materials and Methods) in wild type (*black*; n = 8), *Fgf14^F145S/+^*(*purple*, n = 7), and *Fgf14^F145S/F145S^* (*blue*, n = 7) Purkinje neurons are plotted as a function of the test potential and fitted with first order Boltzmann functions. (**D**). The voltage-dependences of steady-state inactivation of I_NaT_ were also measured in adult *Fgf14^+/-^*(*green*, n = 10) and *Fgf14^-/-^* (*red*, n = 10) Purkinje neurons, plotted and fitted (as in panel **B**), and compared to the steady-state inactivation data for I_NaT_ in wild type (black; n = 11) Purkinje neurons. (**E**). Mean ± SEM normalized peak Nav conductances were also determined in *Fgf14^+/-^* (*green*, n = 7) and *Fgf14^-/-^*(*red*, n = 7) Purkinje neurons and are plotted and fitted (similar to panel **C**) in comparison with data from wild type (*black*; n = 8) Purkinje neurons. (**F**). To compare the kinetics of I_NaT_ decay, the waveforms of the currents evoked at 0 mV in wild type, *Fgf14^-/-^* and *Fgf14^F145S/F145S^* Purkinje neurons were fitted with single exponentials. As is evident, the mean ± SEM time constants determined from these fits indicate that the kinetics of I_NaT_ inactivation are similar in wild type, *Fgf14^-/-^*and *Fgf14^F145S/F145S^* Purkinje neurons.

Similar to Purkinje neurons with targeted knock-in of the *Fgf14^F145S^*mutation, targeted disruptions to one or both *Fgf14* alleles caused significant (*P* ≤ .0001, two-way ANOVA) hyperpolarized shifts in the voltage-dependence of Nav current steady-state inactivation, compared to wild type controls (Figure 4D). Boltzmann fits of normalized Nav current availability (plotted against conditioning voltage) in wild type (n = 11), *Fgf14^+/-^* (n = 10) and *Fgf14^-/-^* (n = 10) Purkinje neurons had V_1/2_ values of -58.0 ± 0.9 mV, -64.0 ± 0.9 mV, and -71.0 ± 1.0 mV, respectively, and slope factor values of 11.3 ± 0.9, 12.1 ± 0.9 and, 10.5 ± 1.0, respectively. Targeted deletion of *Fgf14* did not affect the voltage-dependence of Nav current activation (Figure 4E). The kinetics of I_NaT_ inactivation were measured at a 0 mV in wild type, *Fgf14^F145S/F145S^* and *Fgf14^-/-^* Purkinje neurons by fitting the decay phases of the currents with single exponentials; the mean ± SEM time constant derived from these fits ((Figure 4F) reveal that I_NaT_ inactivation kinetics in *Fgf14^F145S/F145S^* (0.35 ± .02 ms) and *Fgf14^-/-^* (0.31 ± .02 ms) Purkinje neurons are not significantly different from wild type (0.37 ± .03 ms) values.

### Targeted deletion of *Fgf14* does not affect I_NaR_ or the kinetics of I_NaT_ inactivation in acutely isolated neonatal Purkinje neurons

Acute, shRNA-mediated knock-down of iFGF14 in isolated neonatal cerebellar Purkinje neurons was previously reported to accelerate the kinetics of I_NaT_ inactivation and to reduce the density of the resurgent Nav current (I_NaR_) (Yan et al., 2014). As illustrated in Figure 4C, we found that the mean ± SEM time constants of I_NaT_ inactivation in adult *Fgf14^F145S/F145S^* and *Fgf14^-/-^* Purkinje neurons in acute cerebellar slices are similar to wild type controls. To further explore the effects of the loss of iFGF14 on Nav currents in these cells, we obtained voltage-clamp recordings from acutely isolated (Figure 5A) wild type and *Fgf14^-/-^* neonatal (P11-P16) Purkinje neurons and measured I_NaT_ inactivation kinetics and I_NaR_ density. Representative I_NaR_ recorded in wild type and *Fgf14^-/-^* Purkinje neurons are shown in Figure 5B. As is evident, these experiments revealed I_NaR_ measured in wild type and *Fgf14^-/-^*Purkinje neurons are similar in magnitude and voltage-dependences of activation (Figure 5C).These plots are also similar when I_NaR_ is normalized to peak I_NaT_ (measured at 0 mV, Figure 5D) or whole-cell membrane capacitance (Figure 5E). Similar to Fig. 4F, I_NaT_ inactivation kinetics were measured by fitting the decay phase of I_NaT_ to first order exponentials. Compared to isolated wild type Purkinje neurons, the kinetics of I_NaT_ inactivation were also unaffected in *Fgf14^-/-^* isolated Purkinje neurons. Mean ± SEM time constants of I_NaT_ inactivation at 0 mV and -20 mV measured in wild type (0 mV, 0.33 ± .04ms; -20mV, 0.44 ± .05 ms) and *Fgf14^-/-^* (0mV, 0.29 ± .02 ms; -20mV, 0.48 ± .03 ms) Purkinje neurons are plotted in Figure 5F.

**Figure 5.**
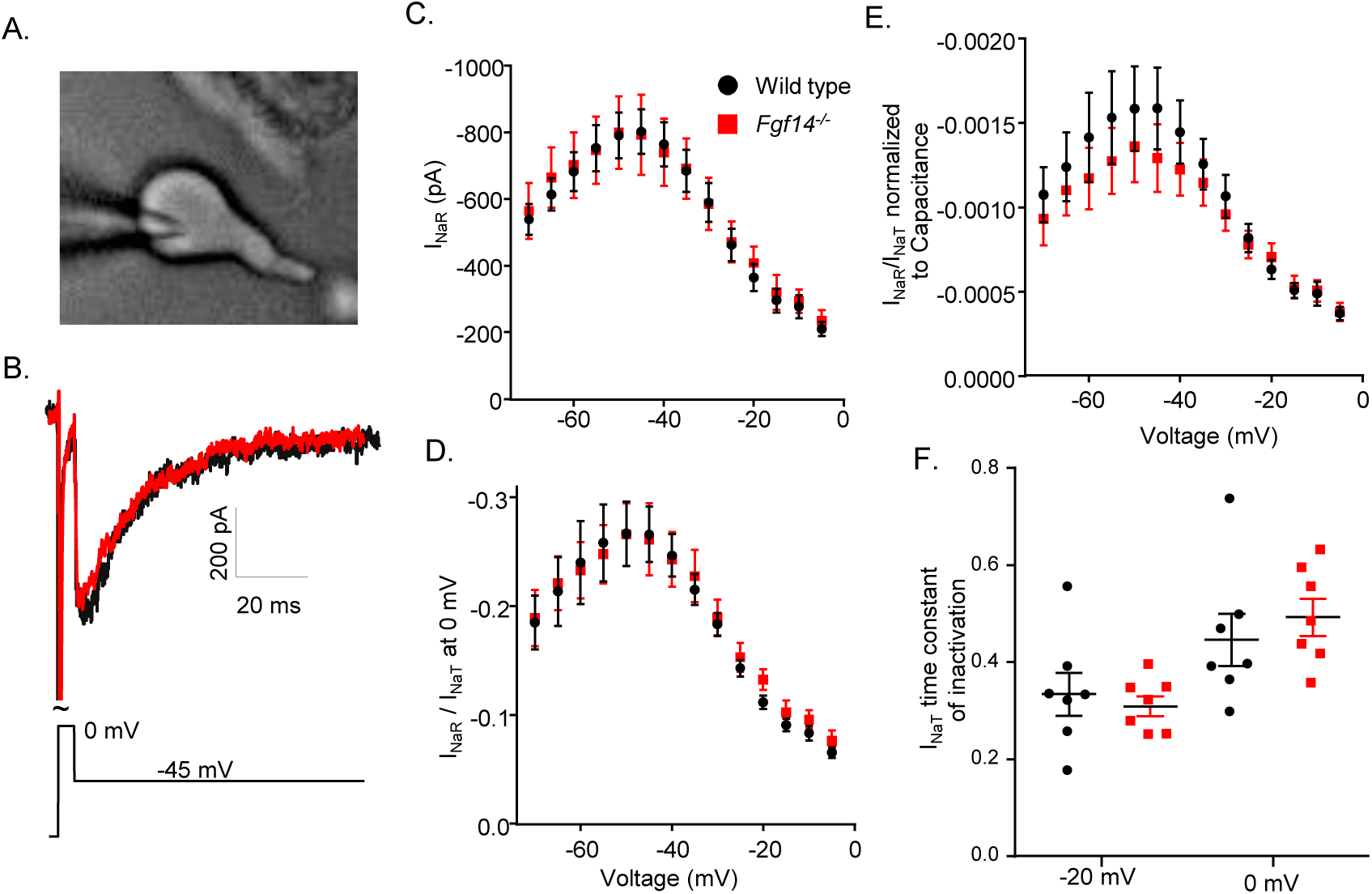
Targeted deletion of *Fgf14* does not affect I_NaR_ or the kinetics of I_NaT_ inactivation in acutely dissociated Purkinje neurons. (**A**). Differential interference contrast (DIC) image of an acutely isolated neonatal cerebellar Purkinje neuron during whole-cell patch clamp recording. (**B**). Representative TTX-sensitive I_NaR_ recorded during a voltage step to -45 mV following a 5 ms depolarization to 0 mV in acutely isolated wild type (*black*) and *Fgf14^-/-^* (*red*) Purkinje neurons with 151 mM Na^+^ in the bath; the voltage command is displayed below the current records. (**C-E**). Mean ± SEM peak I_NaR_ amplitudes (**C**), peak I_NaR_ normalized to peak I_NaT_ at 0 mV (**D**), and I_NaR_/I_NaT_ normalized to cell capacitance (**E**) measured in acutely isolated neonatal *Fgf14^-/-^* (*red*, n = 11) and wild type (*black*, n=15) Purkinje neurons and plotted as a function of voltage, are indistinguishable in magnitude and voltage-dependence. Note that in all cases, I_NaP_, measured 80 ms into the sweep, was digitally subtracted prior to determination of peak I_NaR_. (**F**). The mean ± SEM time constants of I_NaT_ inactivation, measured at -20 mV and 0 mV, are also similar in isolated neonatal wild type (n = 7) and *Fgf14^-/-^*(n = 7) Purkinje neurons.

### *Fgf14^F145S/F145S^* animals have motor deficits similar to *Fgf14^-/-^*animals

The balance and motor coordination of 6-8 week old animals harboring the *Fgf14^F145S^* mutation were examine using the elevated balance beam test (Carter et al., 2001), and compared directly to age-matched animals with targeted deletion of *Fgf14* and wild type controls. A schematic of the elevated balance beam (described in Materials and Methods) is shown in Figure 6A. During five consecutive testing days, adult animals individually traversed a narrow (11 mm wide) beam from a clear platform into an enclosed box. *Fgf14^F145S/F145S^* animals (N = 7) displayed impaired performance, compared to wild type controls, and these deficits, measured as time to cross and the number of hind-limb foot-slips, were similar to deficits measured in *Fgf14^-/-^*(N = 6) animals (Figure 6B, C). Interestingly, *Fgf14^F145S/+^* (N = 7) and *Fgf14^+/-^* (N = 8) animals performed similarly on the elevated balance beam to wild type (N = 8) controls (Figure 6B, C). Older (16- and 32-week-old), *Fgf14^F145S/+^* animals also performed similarly to age-matched wild type animals on the elevated balance beam (data not shown).

**Figure 6.**
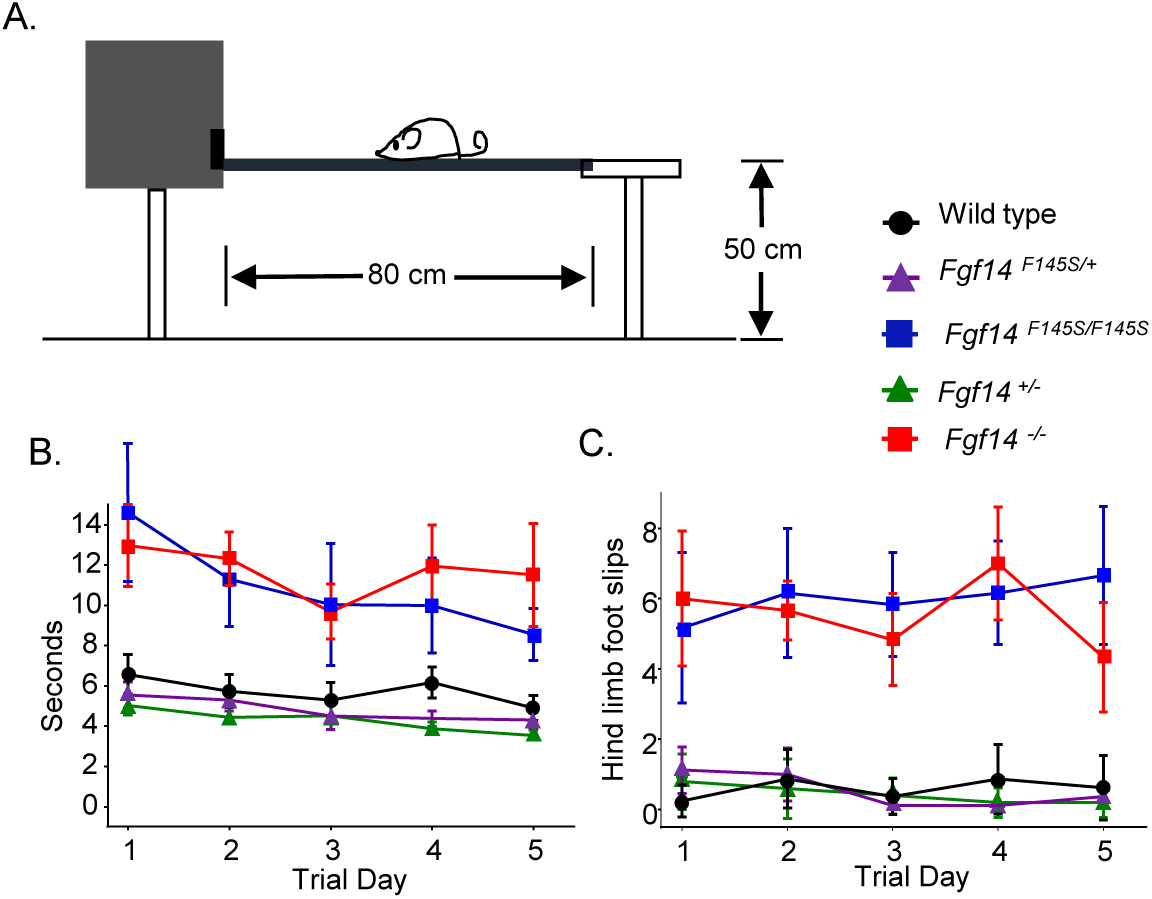
Expression of the *Fgf14^F145S^* mutant allele results in behavioral deficits similar to targeted *Fgf14* deletion. (**A**). Balance and motor coordination were evaluated and compared across genotypes by measuring performance on the elevated balance beam (see Materials and Methods). (**B**). The mean ± SEM times to cross an 11 mm (wide) beam were significantly (*P* < .0001; two-way ANOVA) longer for *Fgf14^F145S/F145S^* (N = 7) and *Fgf14^-/-^* (N = 6) animals, than for wild type (N = 8) and/or heterozygous (*Fgf14^+/-^,* N = 8, and *Fgf14^F145S/+^*, N = 8) animals. (**C**). Analysis of the numbers of hindlimb foot-slips revealed significantly (*P* < .0001; two-way ANOVA) more errors for *Fgf14^F145S/F145S^* and *Fgf14^-/-^* animals, than wild type animals. The performance of heterozygous (*Fgf14^+/-^* and *Fgf14^F145S/+^*) animals was also similar to wild type controls.

### Evoked firing in *Fgf14^F145S^* mutant Purkinje neurons is rescued by membrane hyperpolarization

The results of the voltage-clamp experiments presented in Figures 4 and 5 suggest the impaired excitability of Purkinje neurons with targeted *Fgf14* deletion, or knock-in of the *Fgf14^F145S^* human mutation, is the result of the hyperpolarized shift in the voltage-dependence of steady-state inactivation of I_NaT_. We tested if hyperpolarizing the membranes of these cells, which should drive recovery of inactivated Nav channels, is effective in recovering subsequent evoked firing frequencies. Evoked firing, driven by depolarizing current injections, was measured after a -0.5 nA prepulse (Figure 7A, *right*). The -0.5 nA prepulse silenced spontaneous firing across genotypes and resulted in similar resting membrane potentials (∼-60 mV, Figure 7B), prior to the depolarizing current injections. In Purkinje neurons from wild type animals, mean ± SEM evoked firing frequencies were significantly (*P* ≤ .0001, two-way ANOVA) higher than in Purkinje neurons from *Fgf14^+/-^* and/or *Fgf14^-/-^* animals (Figure 7C), and also higher (*P* ≤ .001, two-way ANOVA) than in Purkinje neurons from *Fgf14^F145S/F145S^* and/or *Fgf14^F145S/+^* animals (Figure 7D). In addition, Purkinje neurons from animals with one wild type *Fgf14* allele (*Fgf14^F145S/+^* and *Fgf14^+/-^*) displayed an intermediate evoked firing phenotype, firing at frequencies that were significantly higher (*P* ≤ .0001, two-way ANOVA) than those measured in *Fgf14^F145S/F145S^*and *Fgf14^-/-^* Purkinje neurons. The intermediate phenotypes in evoked firing rates measured in heterozygous animals are similar to the intermediate shifts in the voltage-dependences of Nav current steady-state inactivation measured in *Fgf14^F145S/+^* and *Fgf14^+/-^* Purkinje neurons (Figure 4). To test the hypothesis that the observed differences in the voltage-dependence of Nav current steady-state inactivation (Figure 4) underlie the reduced firing rates in *Fgf14^+/-^* Purkinje neurons, we hyperpolarized these cells to more negative membrane potentials (i.e., negative to -60 mV), to enable further recovery of inactivated Nav channels. These membrane potentials are plotted as different colors in Figure 7E from two representative *Fgf14^+/-^* cells. As is evident, the results of these experiments indicate that evoked firing frequencies can be increased further if the cells are hyperpolarized to more negative potentials (than -60 mV), consistent with additional recovery of inactivated Nav channels.

**Figure 7.**
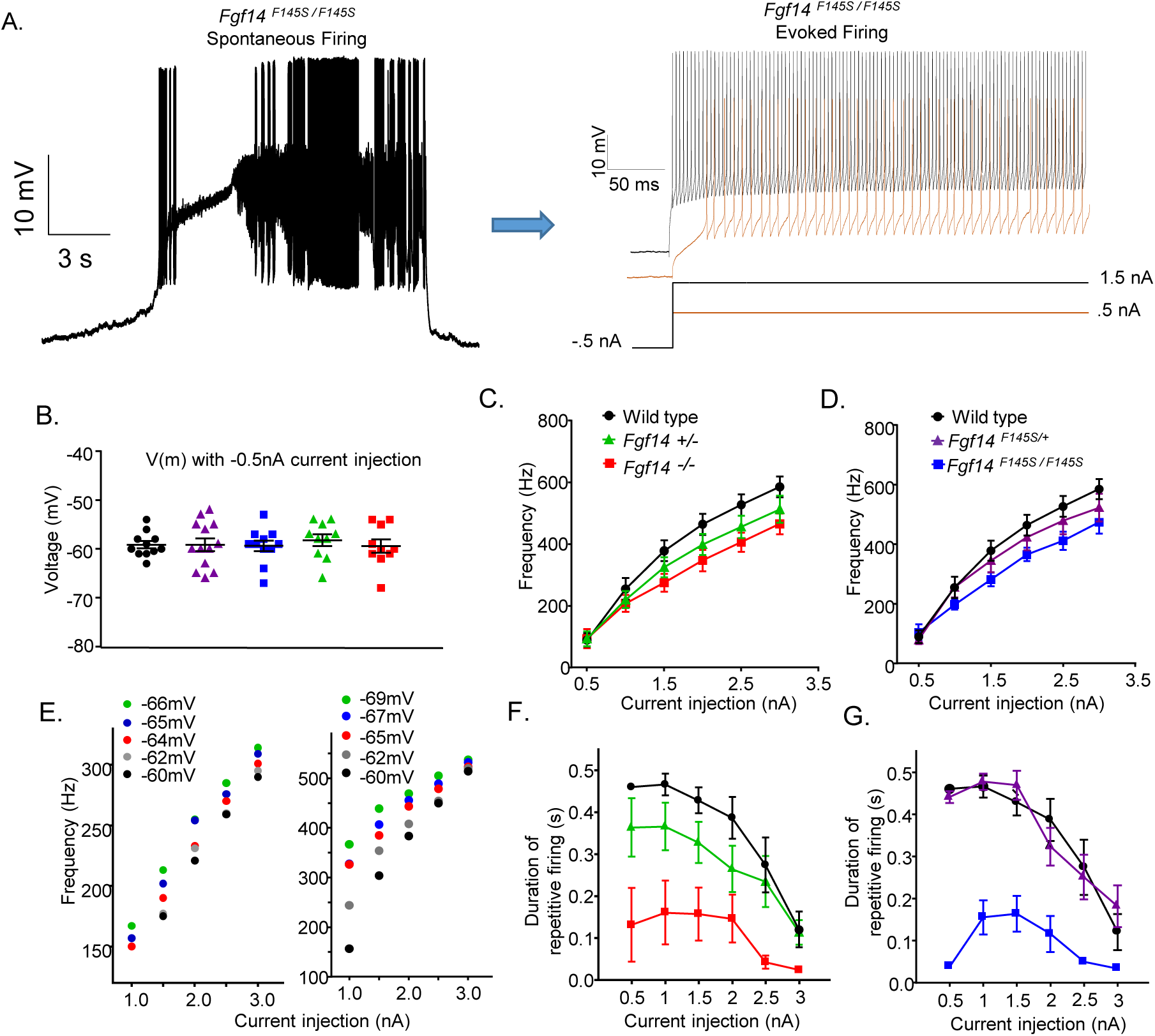
Membrane hyperpolarization rescues firing in *Fgf14^F145S^* mutant and *Fgf14* knock-out Purkinje neurons. (**A**). Representative recordings of firing evoked by 0.5 nA and 1.5 nA current injections in an adult wild type Purkinje neuron are shown. Prior to depolarizing current injections, -0.5 nA current injections were used to hyperpolarize the cell. Current injection protocols are shown below the voltage records. The -0.5 nA pre-pulses resulted in similar hyperpolarizations of the membrane potentials of Purkinje neurons of all genotypes, as illustrated in panel (**B**). (**C-D**). Mean ± SEM evoked firing frequencies of wild type, *Fgf14* knock-out, and *Fgf14^F145S^* knock-in Purkinje neurons are plotted against the amplitudes of depolarizing current injections. As is evident, evoked firing frequencies are similar in heterozygous mutant (*Fgf14^+/-^* and *Fgf14^F145S/+^*) and in homozygous mutant (*Fgf14^-/-^* and *Fgf14^F145S/F145S^*) Purkinje neurons. (**E**). Evoked firing frequencies are plotted against the amplitudes of the depolarizing current injections presented after hyperpolarizing prepulses to varying voltages (plotted in different colors) in two representative *Fgf14^+/-^* cells. In both *Fgf14^+/-^* Purkinje neurons, increasing the amplitude of the membrane hyperpolarization (prior to the depolarizing current injection) resulted in higher evoked firing frequencies. (**F-G**). Durations of repetitive firing during the 0.5 s current injection in wild type, *Fgf14* knock-out, and *Fgf14^F145S^* knock-in adult Purkinje neurons are plotted against the depolarizing current injection. Similar levels of depolarization block is observed in Purkinje neurons from heterozygous (*Fgf14^+/-^*and *Fgf14^F145S/+^*) and homozygous (*Fgf14^-/-^* and *Fgf14^F145S/F145S^*) animals.

The rescue of evoked firing (after hyperpolarization) in Purkinje neurons lacking *Fgf14* is expected to be temporary during depolarizing current injections because, during the sustained depolarizing current injection, Nav channels should again accumulate into an inactivated state. We measured the rate at which Purkinje neurons enter a state of depolarization block during the 0.5 s depolarizing current injections. The mean ± SEM firing duration is plotted against the depolarizing current injection in wild type, *FGF14^F145S^*knock-in, and *Fgf14* knock-out Purkinje neurons in Figure 7F and 7G. These experiments revealed that *Fgf14^-/-^* and *Fgf14^F145S/F145S^*Purkinje neurons undergo depolarization block more rapidly (*P* ≤ .0001, two-way ANOVA) than wild type, *Fgf14^+/-^* (Figure 7F) and/or *Fgf14^F145S/+^*(Figure 7G) Purkinje neurons.

## Discussion

### *Fgf14^145S^* is a Loss-of-function Mutation

We developed a knock-in mouse model to investigate the pathophysiology underlying the disease phenotypes caused by the *FGF14^F145S^* human mutation. Electrophysiological analyses revealed that Purkinje neurons from adult *Fgf14^F145S/+^* and *Fgf14^+/-^* animals lack the ability to sustain continuous repetitive firing, which is a hallmark feature of wild type Purkinje neurons. Voltage-clamp experiments revealed the observed deficits in repetitive firing are caused by shifts (in the hyperpolarized direction) of the voltage-dependence of I_NaT_ inactivation in *Fgf14^F145S/+^* and *Fgf14^+/-^* Purkinje neurons. These hyperpolarized shifts were intermediate to the shifts measured in *Fgf14^F145S/F145S^*and *Fgf14^-/-^* Purkinje neurons. Together, these experiments indicate the *FGF14^145S^* mutation causes loss of iFgf14 function and haploinsufficiency in *Fgf14^F145S/+^* Purkinje neurons, which result in phenotypes that are similar to those found in *Fgf14^+/-^* Purkinje neurons. Indeed, animals expressing two mutant alleles (*Fgf14^F145S/F145S^*) phenocopy *Fgf14^-/-^* animals and Western blot results indicate the iFGF14^F145S^ mutant protein is not detected in cerebellar tissue lysates taken from adult *Fgf14^F145S/F145S^* animals. In *Fgf14^F145S/+^* cerebellar tissue lysates, the anti-iFGF14 signal is comparable to *Fgf14^+/-^* animals and is reduced from wild type levels suggesting that the mutant protein likely is not stable and is, therefore, degraded. Interestingly, in the original description of the *FGF14^F145S^*human mutation, van Swieten et al. (2003) noted that the amino acid change from a phenylalanine to a serine at position 145, which lies in the central core of iFGF14, would be expected to reduce the stability of the iFGF14 protein, resulting in its degradation.

Partial block of Nav channels using a low concentration (1 nM) of TTX produced firing phenotypes that are similar to those observed in *Fgf14^+/-^* and *Fgf14^F145S/+^* Purkinje neurons (Figure 2), supporting the hypothesis that a loss of available Nav channels is responsible for the altered firing phenotypes of *Fgf14^+/-^*and *Fgf14^F145S/+^* Purkinje neurons. Hyperpolarizations of the resting membrane potential of *Fgf14^+/-^* Purkinje neurons (prior to depolarizing current injections), which allows for the recovery of inactivated Nav channels, resulted in increased evoked firing rates in *Fgf14^+/-^*Purkinje neurons (Figure 7E). Previously, it was reported that shRNA-mediated knock-down of iFGF14 in cultured Purkinje neurons significantly accelerates the inactivation kinetics of I_NaT_ and reduces the levels of I_NaR_ when compared to cultured Purkinje neurons expressing a scrambled control shRNA (Yan et al., 2014). In the presence of TTX and with reduced (50 mM) extracellular sodium, White et al. (2019) also reported accelerated I_NaT_ inactivation and reduced I_NaR_ in acutely isolated *Fgf14^-/-^* Purkinje neurons. Under physiological (151 mM) extracellular sodium, however, we found that the kinetics of I_NaT_ inactivation, as well as the voltage-dependence and the magnitude of I_NaR_, are similar in acutely isolated *Fgf14^-/-^*and wild type Purkinje neurons (Figure 5B-D). Together, these results support the hypothesis that hyperpolarized shifts in the voltage-dependence of I_NaT_ inactivation are responsible for the reduced firing in Purkinje neurons that lack two wild type *Fgf14* alleles.

Interestingly, Laezza et al. (2007) showed that transfection of cDNA constructs encoding iFGF14^F145S^ in cultured hippocampal neurons disrupts the interaction between wild-type iFGF14 and Nav α subunits, and affects firing in these cells via a dominant-negative mechanism. Yan et al. (2013) also showed that the acute expression of iFGF14^F145S^ in cultured granule neurons causes a dominant-negative reduction of Ca^2+^ currents and vesicular recycling. These previous experiments and the results presented here indicate that there are clear and important differences in the effects of acutely expressed *FGF14^F145S^* cDNA constructs in cultured neurons and the effects of genomic *F14^F145S^* expression, measured in acute preparations. One possible explanation is that the transfection of cultured neurons with *FGF14^F145S^* containing plasmids, driven by the CMV promoter, causes gross over-expression of the mutant protein that results in dominant-negative effects that do not occur under native iFGF14^F145S^ expression conditions.

Adult (6-8 week-old) animals that express one wild type *Fgf14* allele (*Fgf14^+/-^* and *Fgf14^F145S/+^*) performed similarly to age-matched wild type controls on the elevated-balance beam test, indicating that mice harboring a single *Fgf14^F145S^* mutation do not recapitulate the motor phenotypes expressed in humans. Conversely, *Fgf14^-/-^*and *Fgf14^F145S/F145S^* animals had severe motor deficits at 6-8 weeks-old. The absence of a motor phenotype in *Fgf14^F145S/+^* and *Fgf14^+/-^* animals is surprising, given that there are clear deficits in the intrinsic firing properties of Purkinje neurons in acute brain slices from adult *Fgf14^+/-^* and *Fgf14^F145S/+^*animals. One possibility is that the preparation of acute sagittal brain sections, and the disruption of physiological synaptic- and neuromodulatory-influences on Purkinje neuron firing, exposes differences in *Fgf14^+/-^*and *Fgf14^F145S/+^* Purkinje neuron membrane excitability, when compared to wild type Purkinje neurons; however, in the intact cerebellar circuits, these changes in membrane excitability are insufficient to disturb cerebellar function. *In vivo* electrophysiological experiments will be necessary to test this possibility directly.

## Acknowledgments

The authors thank Drs. A. Burkhalter and T. Hermanstyne for many helpful discussions. We also thank R. Wilson, R. Mellor and C. Smith for expert technical assistance. Financial support provided by the NIH (R01NS065761 to JMN and DMO, F32NS090765 to JLR, R15NS125560 to JLR).

## Figure Legends

**Extended Data Figure 3-1.**

Full Western blot of fractionated total protein lysates, prepared from cerebellar tissues isolated from adult wild type, *Fgf14^F145S/+^*, *Fgf14^F145S/F145S^*, *Fgf14^+/-^*, and *Fgf14^-/-^*animals, probed with a rabbit polyclonal α-iFGF14 antiserum (Rb-α-iFGF14), are shown; the arrowheads indicate the iFGF14 protein (reduced in the *Fgf14^F145S/+^*and *Fgf14^+/-^* samples and missing in the *Fgf14^F145S/F145S^* and *Fgf14^-/-^* samples). As shown in the right panel, membranes were also probed with a mAb-α-tubulin antibody. The regions outlined by the blue boxes indicate the parts of the blots shown in Figure 3E.

## Notes

**Conflict of interest statement:** The authors have no conflicting interests to declare.

### Competing Interest Statement

The authors have declared no competing interest.

