## Supplementary figures and images for "*In Vivo* Expression of an SCA27A-linked *FGF14* Mutation Results in Haploinsufficiency and Impaired Firing of Cerebellar Purkinje Neurons"

### Extended Data Figure 3-1

#### Extended Data Figure 3-1

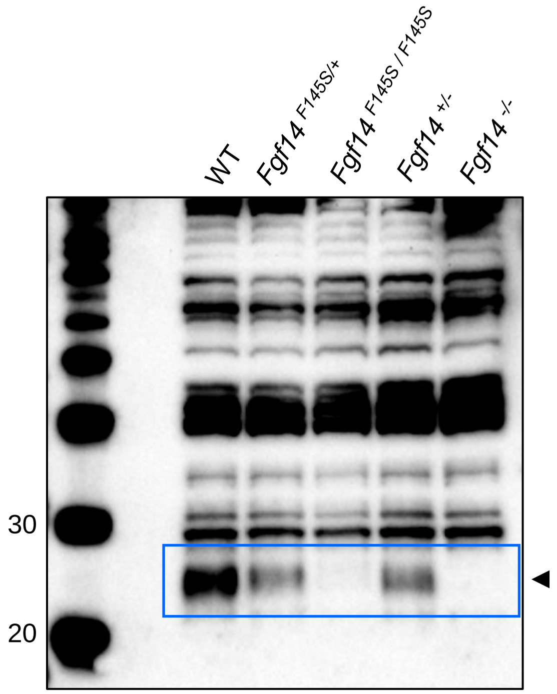


Rb-α-iFGF14

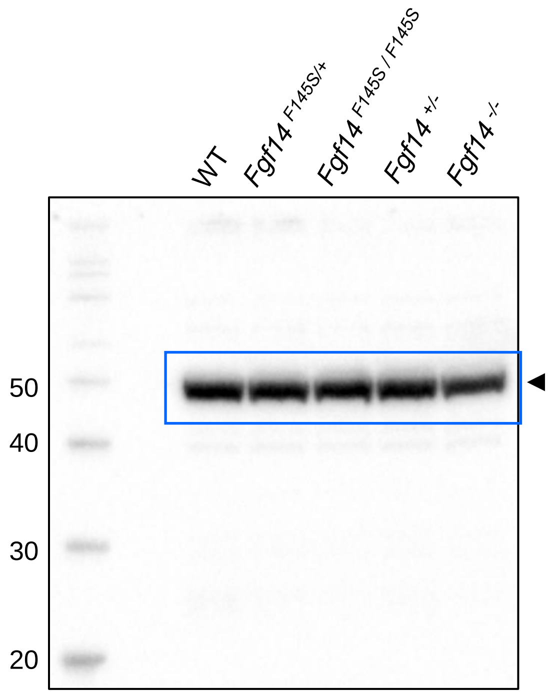


mAb-α-Tubulin
